# A Genome-Wide Association Analysis of Happiness: Consistent Genetic Effects Across the Lifespan and Across Genetic Ancestries in Multiple Cohorts

**DOI:** 10.1101/2022.04.05.487098

**Authors:** Joey Ward, Laura Lyall, Breda Cullen, Rona J. Strawbridge, Xingxing Zhu, Ioana Stanciu, Alisha Aman, Claire L. Niedzwiedz, Jana Anderson, Mark E. S. Bailey, Donald M. Lyall, Jill Pell

## Abstract

We present a genome-wide association study of a general happiness measure in 118,851 participants from the UK Biobank. Using BOLT-LMM, we identify 3 significant loci with a heritability estimate of 0.8%. Linkage disequilibrium score regression was performed on the ‘big five’ personality traits finding significant associations with lower neuroticism and higher extraversion and conscientiousness. Using a novel approach, we construct LDpred-inf polygenic risk scores in the Adolescent Brain Cognitive Development (ABCD) cohort and the Add Health cohort. We detected nominally significant associations with several well-being measures in ABCD and significant correlations with a happiness measure in Add Health. Additionally, we tested for associations with several brain regions in a white British subsample of UK Biobank finding significant associations with several brain structure and integrity phenotypes.

We demonstrated a genetic basis for general happiness level and brain structure that appears to remain consistent throughout the lifespan and across multiple ancestral backgrounds.

**Author summary:** At the genetic level, there has been little investigation into whether people may have a baseline happiness level which varies from person to person. Here we perform a genetic analysis in the UK Biobank to identify three genetic loci that associate with general happiness level and preform genetic correlations of our results with the ‘Big Five’ personality traits, identifying significant correlations with neuroticism, conscientiousness and extraversion.

We use the resulting summary statistics to create LDpred-inf polygenic risk scores in UK biobank identifying several brain metrics and regions associate with genetic loading for general happiness level. We also use a novel method to create LDpred-inf polygenic risk scores in two other cohorts, ABCD and Add Health. We found significant correlations with an independent happiness measure in Add Health and nominally significant correlations with several well-being measures in ABCD in both those of European Ancestry and all other ancestries found in these cohorts. We also attempted to replicate our UK Biobank MRi finding in ABCD.

We conclude there is evidence that individuals have a general happiness level that is in part genetic which spans across age and ancestry.

## Introduction

Happiness is the core positive emotional state. As an emotional trait that is affected in clinical outcomes (e.g. lack of happiness in depression and exacerbation of happiness in the manic phase of bipolar disorder) it can be considered a positive valence Research Domain Criteria (RDOC) trait^1^. At the genetic level it is usually analysed as part of a wider concept of mental well-being^2^. There is evidence that, generally, an individual has a baseline happiness level which remains relatively stable over time^3^ even after major positive or negative life events such as winning the lottery or being the victim of an accident^4^.

There has been little investigation into the genetics of general happiness level and most of the research has been performed via twin studies^5^, with a recent meta-analysis giving an estimate for heritability of a ‘well-being’ trait of 36% and a slightly lower estimate of 32% for heritability of ‘life satisfaction’ ^6^. However, to date there has been no genome-wide association study (GWAS) of a general happiness measure to establish features of its genetic architecture and to identify specific areas of the genome that contribute to this trait. Greater understanding of the genetic basis of subjective happiness could be useful in identifying whether and how targeted interventions could improve happiness, both in the general population and in clinical populations^5^. This in turn will increase our understanding of psychology and neurodevelopment aiding in the future treatment of patients^7^.

Here we report the first GWAS of general happiness in individuals of white British ancestry, as self-reported in the UK Biobank^8^ cohort, using BOLT-LMM^9^ and the use of polygenic risk scores (PRS) in two additional cohorts: the Adolescent Brain Cognitive Development (ABCD) cohort^10^ and the National Longitudinal Study of Adolescent to Adult Health (Add Health)^11^. This study had three aims: (1) to identify specific loci in the genome that are associated with self-reported general happiness level; (2) to test whether increased genetic loading of this measure is significantly associated with happiness and well-being measures in independent cohorts that span differing age ranges and different ancestries compared to the discovery GWAS cohort; and (3) to investigate whether genetic predisposition for general happiness level is associated with average differences in brain structure.

## Results

### Phenotype stability

We established stability of general happiness level in UK Biobank over time in those with a repeat measure (Pearson’s r = 0.58, S.E. = 0.01, p < 2.2 × 10^−16^, n = 4,703).

Weighted Pearson’s correlations were calculated to establish the stability of the Add Health happiness phenotype using the function weightedCorr of the R package wCorr, which showed a correlation between waves I and II of 0.34, S.E. = 0.008, p < 0.0001 (weighted by wave 2 variable gswgt2) and between waves II and IV of 0.22, S.E. = 0.01, p < 0.0001 (weighted by wave 4 variable gswgt4).

### GWAS in UK Biobank

After all exclusions, the final sample size of the GWAS was 118,851, main due to low response rate of the question. The sample had a mean age of 57.5 (S.D. = 8) years and 53.9% were female. A full breakdown of the demographics at each happiness response level can be found in Table S1.

Three loci that achieved genome-wide significance (Table 1, Figure 1, Figures S1-3). They were rs3129791 on chromosome six, rs72743916 on chromosome nine and rs11474549 on chromosome twenty. The heritability estimate h_2_^SNP^ was 0.0084 (S.E. = 0.004) and there was minimal inflation of the test statistics as λ_GC_ = 1.001. The LDSR intercept was 0.999 (S.E. = 0.004) showing there was no inflation of the test statistics due to unaccounted population stratification.

**Table 1.**
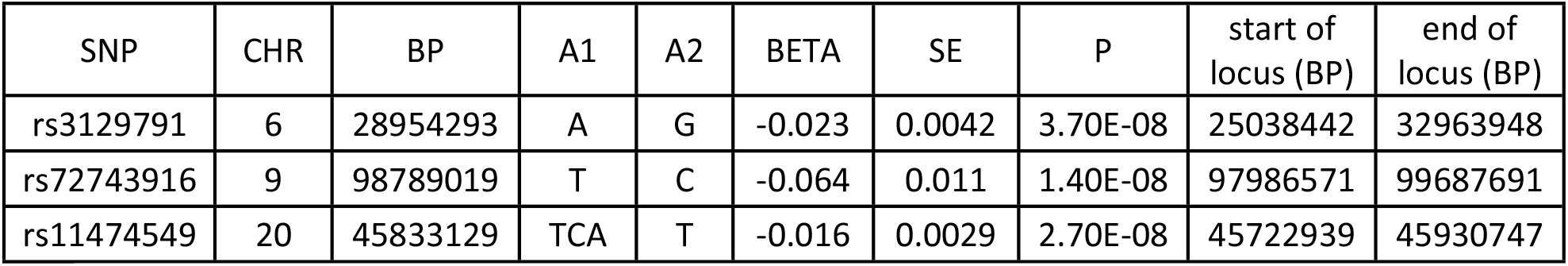
Genomic loci significantly associated with general happiness level.

**Table 1.** Linkage disequilibrium score regression with general happiness level and the ‘Big Five’ personality traits.

| Trait | $r_g$ | se | z | p | P.FDR |
| --- | --- | --- | --- | --- | --- |
| conscientiousness | 0.589 | 0.1549 | 3.80 | 1.43E-04 | 7.15E-04 |
| neuroticism | -0.8448 | 0.33725 | -2.51 | 1.22E-02 | 0.021 |
| extraversion | 0.8025 | 0.32086 | 2.50 | 1.24E-02 | 0.021 |
| openness | -0.1909 | 0.10298 | -1.85 | 6.38E-02 | 0.080 |
| agreeableness | 0.8574 | 0.9374 | 0.91 | 3.60E-01 | 0.360 |

**Figure 1.**
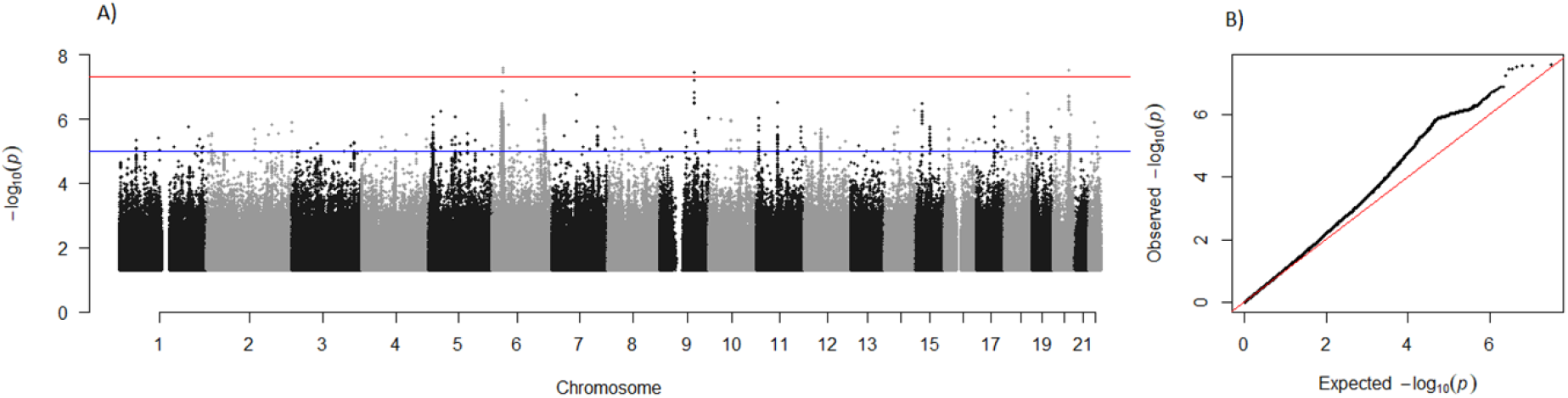
Manhattan plot and QQ plot of general happiness. A) X axis = chromosome from 1 -22 y axis = -log10 p value. Blue line shows suggestive genome wide significance threshold, red line shows genome wide significance threshold. B) x axis = - log10 of the expected p values, y axis = -log10 of the observed p values.

### Linkage Disequilibrium Score Regression

Three of the five tested traits were significant after FDR correction (Table 2). The strongest correlation was with neuroticism which showed an inverse relationship with happiness (rg = -0.84, S.E. = 0.34, p.fdr = 0.021). Both extraversion and conscientiousness showed positive correlations with happiness (extraversion: rg = 0.8, S.E. = 0.32, p.fdr = 0.021, conscientiousness: rg = 0.58, S.E. = 0.16, p.fdr = 0.0007).

**Table 2.** Association of LDpred-inf PRS of general happiness level and well-being measures in ABCD white subsample

| outcome | BETA | S.E | P | P.FDR |
| --- | --- | --- | --- | --- |
| attentive | 8.13E-03 | 0.017 | 0.63 | 0.67 |
| delighted | 0.010 | 0.015 | 0.51 | 0.67 |
| calm | 0.008 | 0.014 | 0.57 | 0.667 |
| relaxed | 0.000 | 0.014 | 0.99 | 0.99 |
| enthusiastic | 0.014 | 0.016 | 0.39 | 0.585 |
| interested | 0.031 | 0.016 | 0.05 | 0.13 |
| confident | 0.027 | 0.015 | 0.07 | 0.15 |
| energetic | 0.009 | 0.016 | 0.55 | 0.679 |
| concentrate | 0.031 | 0.015 | 0.04 | 0.12 |

### Polygenic risk scoring

#### UK Biobank

In those who were excluded from the GWAS due to them having MRI data but who responded to the happiness question and passed genetic quality control (n = 18,795), we detected a significant association of happiness with the LDpred polygenic risk scores (β = 0.057, S.E. = 0.005, p = 4.85*10^−31^) with the model explaining 0.018% of the variance.

#### ABCD

In the white subsample, two of the well-being outcomes tested were significantly correlated with the LDpred happiness PRS (Table 3) but neither survived FDR correction. The strongest associations were with being interested (β = 0.0311, S.E. = 0.016, p = 0.05, p.FDR = 0.13) and being able to concentrate (β = 0.0307, S.E. = 0.015, p = 0.04, p.FDR = 0.12). All of the models showed that greater genetic loading for happiness was associated with greater agreement with the well-being items.

**Table 3.** Association of LDpred-inf PRS of general happiness level and well-being measures in ABCD multi ancestry sample

| outcome | BETA | S.E | P | P.FDR |
| --- | --- | --- | --- | --- |
| confident | 0.030 | 0.013 | 0.018 | 0.0702 |
| enthusiastic | 0.029 | 0.014 | 0.039 | 0.0702 |
| energetic | 0.029 | 0.014 | 0.039 | 0.0702 |
| concentrate | 0.028 | 0.013 | 0.028 | 0.0702 |
| calm | 0.025 | 0.012 | 0.035 | 0.0702 |
| delighted | 0.024 | 0.013 | 0.074 | 0.111 |
| attentive | 0.022 | 0.015 | 0.148 | 0.19 |
| interested | 0.018 | 0.014 | 0.189 | 0.213 |
| relaxed | 0.006 | 0.012 | 0.617 | 0.617 |

In the whole sample, five of the nine aspects of well-being tested were nominally significant; however, none survived FDR correction (Table 4). The strongest association was with being confident (β = 0.03, S.E. = 0.013, p = 0.018, p.FDR = 0.07). Being enthusiastic and energetic showed the same strength of effects (both: β = 0.029, S.E. = 0.014, p = 0.039, p.FDR = 0.07) followed by being able to concentrate (β = 0.028, S.E. = 0.013, p = 0.028, p.FDR = 0.07) and being calm (β = 0.025, S.E. = 0.012, p = 0.035, p.FDR = 0.07). As with the white subsample analysis, all the models showed that greater genetic loading for happiness associated with greater agreement with the well-being items.

The results of the ABCD P&T models generally reflected those of the LDpred scores (Tables S2 and S3). In the white subsample, being interested and able to concentrate were nominally significant. In the P&T scores, being interested was significant at the p < 0.05 threshold and showed the same direction of effect as did all the other interested outcome models. Similarly, with concentration, models were significant at the p < 0.01 threshold and higher and all models showed the same direction of effect as the LDpred model. Two differences in the P&T models compared to the LDpred models were that being delighted was significant at the p < 5*10^−5^ threshold (β = 0.039, S.E. = 0.016, p = 0.013) and being confident was significant at the p < 0.05 (β = 0.038, S.E. = 0.015, p = 0.014) and p = 1 thresholds (β = 0.034, S.E. = 0.016, p = 0.027). All other P&T models in the white subsample did not achieve statistical significance.

In the whole sample, being confident, enthusiastic, energetic, able to concentrate and calm were all nominally significant. In the P&T models, being confident and being enthusiastic were both significant at the p < 0.05 threshold (confident: β = 0.028, S.E. = 0.013, p = 0.029. enthusiastic: β = 0.027, S.E. = 0.014, p = 0.05). Being energetic was nominally significant at p <0.5 (β = 0.021, S.E. = 0.009, p = 0.017) and p = 1 thresholds (β = 0.02, S.E. = 0.086, p = 0.02).\ Being able to concentrate showed greatest concurrence with all models being nominally significant from p < 0.01 and above. This outcome showed the strongest association in the LDpred models and had the most significant thresholds in the P&T models. Being calm was significant at both the p < 0.5 (β = 0.024, S.E. = 0.02, p = 0.04) and the p = 1 (β = 0.025, S.E. = 0.02, p = 0.037) thresholds. All the other P&T models did not achieve statistical significance. Also note that all P&T models like the LDpred models would not pass FDR correction.

#### Add Health

A significant association was found in the LDpred analysis in the European ancestry subsample (β = 0.036, S.E. = 0.012, p = 0.0019) and also in the whole sample (β = 0.025, S.E. = 0.009, p = 0.009). The P&T models reflected similar findings with significant associations found in the European ancestry subsample (best threshold p <0.05; β =0.03, S.E. = 0.012, p = 0.01), and in the whole sample (best threshold p <0.1; β =0.022, S.E. = 0.009, p = 0.017) (Tables S4 and S5 respectively). In both the European subsample and in the whole sample the models showed a positive correlation with greater genetic loading for happiness with the exception of the p < 5 *10^−5^ threshold in both groups and p < 0.01 threshold in the whole sample however, none of these models were statistically significant.

### MRI

#### UK Biobank

After correcting for multiple testing via FDR, both (head-size adjusted) white and grey matter volumes (white β = 0.014, S.E. = 0.0053, p.fdr = 0.04; grey β = 0.012, S.E. = 0.0042, p.fdr = 0.016), as well as the general factor for frontal lobe volume (β = 0.019, S.E. = 0.0048, p.fdr = 0.001), remained significant, showing that increasing genetic loading for happiness is associated with higher values of these brain measures (Table S6). We also detected significant associations between genetic loading for general happiness and the left hippocampus tail (β =0.015, S.E. = 0.005, p.fdr = 0.013) and nominally significant associations with the right hippocampus body (β =0.012, S.E. = 0.005, p = 0.021, p.fdr = 0.078) and the left and right thalamus (left: β =0.011, S.E. = 0.0048, p = 0.024, p.fdr = 0.078. right: β =0.01, S.E. = 0.0048, p = 0.038, p.fdr = 0.11).

The DTI MRI parameters MD and FA both associated with genetic loading for happiness. Greater FA (reflecting better white matter integrity) was associated with higher genetic loading for happiness while the effect was in the opposite direction for MD, for which higher values reflect worse white matter integrity (FA: β = 0.019, S.E. = 0.0056, p.fdr = 0.009; MD: β = -0.021, S.E. = 0.0054, p.fdr < 0.001)

#### ABCD

In the whole sample (Tables S7) greater right amygdala volume (β = 0.031, S.E. = 0.01, p = 0.005, p.fdr = 0.1), greater white matter volume (β = 0.028, S.E. = 0.01, p = 0.01, p.fdr = 0.1) and greater frontal lobe volume (β = 0.023, S.E. = 0.01, p = 0.035, p.fdr = 0.2) were all nominally significantly associated with increased genetic loading for general happiness, although none passed FDR correction. Both MD and FA, which were significant in UK Biobank, as well as left and right thalamus, which were nominally significant in the UK Biobank, had the same direction of effect in the whole sample ABCD models. The same was true for all the directions of effect in the white sub sample but none of these models were significant (Table S8).

## Discussion

We have shown that a single-item, self-reported happiness measure can be used to identify loci in the genome that are associated with happiness, adding to the evidence that individuals do have a baseline happiness level. Given that our sample size was moderate, the heritability estimate was far below that determined by the twin study meta-analysis of Bartels^6^. The mechanism by which the genes at these loci may be affecting the phenotype is unclear; each locus identified in the GWAS does contain genes thought to play a role in neurodevelopment, although this may reflect pleiotropy. The major histocompatibility complex (MHC) region on chromosome 6 contains hundreds of genes in very high LD. Genes in the MHC region have been shown to be involved in several aspects of neural development, including neurite outgrowth and synapse formation^12^. The most statistically significant SNP within the locus identified on chromosome 9 was rs72743916, located in an intron of *ERCC6L2*. Mutations in this gene have been shown to cause bone marrow failure syndrome 2, symptoms of which include developmental delay and learning difficulties^13^. The third locus, identified on chromosome 20, is found between *EYA2* and *ZMYND8*. No mutations in *EYA2* have been associated with Mendelian disorders, but over-expression has been found in astrocytoma (a type of brain tumour)^14^. ZMYND8 is a transcriptional regulator^15^ and has been shown to bind Drebrin which regulates the distribution of ZMYND8 protein between nucleus and the synapse^16^.

There were significant genetic correlations between the happiness phenotype and three of the ‘big five’ personality traits tested and the direction of these effects is generally in agreement with the psychological trait literature. Our results were consistent with a meta-analysis of phenotypic happiness and the big five by Steel et al.^17^ as well as work by Hayes and Joseph^18^ showing that conscientiousness and extraversion positively correlate with general happiness whereas neuroticism shows a strong negative correlation. The genetic correlation with openness matched that of Hayes and Joseph in that it was negative (though non-significant), whereas Steel et al. showed a significant positive correlation. With agreeableness our results reflected that of Steel et al in that the correlation was positive (though non-significant) whereas Hayes & Joseph showed a non-significant negative correlation.

The results of the LDpred PRS analyses in ABCD and Add Health showed that the genetic loading of this happiness measure not only remained consistent across the lifespan from age 12 to 73 in those of European ancestry but also across multiple ancestral backgrounds giving evidence that the pathways involved are common to a range of ancestral backgrounds. Although the models for the ABCD cohort were not significant there was a consistent direction of association with all nine of the tested traits in both the white subsample and the multi-ancestry analyses. These models may have been limited by relatively small sample size and also in that the measures used reflect the responses of the parent and not of the child themselves. They are also measures of well-being and not a direct happiness measure.

The use of a novel method involving use of UK Biobank participants as the discovery set for the construction of LDpred scores was also validated by similar results in the P&T PRS that used participants of the respective cohorts to establish LD. This allowed for the sample size to be maximised in these smaller cohorts increasing the statistical power of the analyses.

The results of the MRI analyses detected several brain regions associated with genetic loading for happiness. The frontal lobe has been implicated in hedonic emotions^19,20^ and we showed that greater values of a general factor for frontal lobe volume significantly correlated with genetic loading for happiness; however we were unable to replicate this finding in the ABCD cohort which may be due to a lack of power as the sample size was only approximately 7000. It is also the case that the frontal lobes continue to develop until around age 25 years so ABCD participants may yet be too young for this relationship to be demonstrated. We also detected a positive correlation with white and grey matter volumes in the UK Biobank, which was reflected in the ABCD analyses but only white matter in the whole sample achieved nominal statistical significance.

Larger left hippocampal tail and right hippocampal body both associated with greater genetic loading for happiness whereas the whole hippocampal volume was not significantly associated in either UK Biobank or ABCD. The hippocampus outcomes in both cohorts were insignificant which contradicts what has been found elsewhere in the literature, for example in the meta-analysis of Tanzer & Wayandt^21^, however, they included parahippocampal regions along with the hippocampus. There were no data available for sub-hippocampal structures in the ABCD so a direct comparison was not possible.

We also found significant correlations with MD and FA showing higher white matter integrity correlated with greater loading for happiness. This result was reflected in the ABCD cohort but did not reach statistical significance.

## Strengths and Limitations

The initial GWAS was large enough to detect significant loci and the use of BOLT-LMM allowed for maximisation of the sample size by accounting for relatedness using the genetic relationship matrix. The GWAS was limited by only including those of white British ancestry. This was due to the imputation panels used in the UK Biobank: both the HRC and the 1000genomes reference panel were European only. Future analyses that include those of other ancestries will be possible once the TOPMED dataset becomes available which has participants imputed using the most appropriate ancestral reference panel. UK Biobank is not representative of the UK general population in that participants are generally healthier and have a higher socioeconomic status than the general population and therefore may have a different happiness level distribution than the UK as a whole^22^. The MRI sample is even less representative in that participation was slightly biased towards the fitter, healthier participants in UK Biobank ^23^.

A similar issue arises in the PRS analyses in that those of European ancestry were used to establish LD structure of the genome. This was due to the lower numbers of the non-European ancestry participants in these cohorts.

## Conclusions

These analyses demonstrate that general happiness level has a genetic contribution which has a consistent effect across age groups and ancestral backgrounds. These analyses not only help increase our understanding of psychology and neurodevelopment, the novel methodology of using UK Biobank participants as a reference panel for LDpred risk scores could be adapted for a wide range of other phenotypes.

It is important to note however, the difference in heritability estimates between our genetic study and those obtained at the phenotypic level suggests that there are still many more genetic loci yet to be discovered that will contribute to baseline happiness level. As such, further analyses will be required using larger datasets with less bias towards those of European ancestry.

## Methods

### Cohorts, genotyping and phenotyping

#### UK Biobank

*Cohort Description*

UK Biobank is a cohort of over half a million UK residents, aged from approximately 40 to 70 years at baseline. It was created to study environmental, lifestyle and genetic factors in middle and older age^8^. Baseline assessments occurred over a 4-year period, from 2006 to 2010, across 22 UK centres. These assessments were comprehensive and included social, cognitive, lifestyle and physical health measures.

UK Biobank obtained informed consent from all participants, and this study was conducted under generic approval from the NHS National Research Ethics Service (approval letter dated 29 June 2021, Ref 21/NW/0157) and under UK Biobank approvals for application #6553 ‘Genome-wide association studies of mental health’ (PI Rona Strawbridge; GWAS) and #17689 (PI Donald Lyall; imaging).

#### Genotyping

In March 2018, UK Biobank released genetic data for 487,409 individuals, genotyped using the Affymetrix UK BiLEVE Axiom or the Affymetrix UK Biobank Axiom arrays (Santa Clara, CA, USA) containing over 95% common content. Pre-imputation quality control, imputation and post-imputation cleaning were conducted centrally by UK Biobank (described in the UK Biobank release documentation)^24^.

#### Phenotyping and exclusion criteria

The GWAS was based on the response to the question “In general, how happy are you?” (Data Field 4526), with ordinal responses on a six-point scale ranging from extremely happy to extremely unhappy. Participants were excluded if they were missing more than 10% of their genetic data, if their self-reported sex did not match their genetic sex, if they were determined by UK Biobank to be heterozygosity outliers, and if they were not of white British ancestry (classified by UK Biobank based on self-report and genetic principal components)^24^. Those who had attended for magnetic resonance imaging (MRI) were excluded from the GWAS, in order to maintain independence for subsequent analyses of the MRI phenotypes.

#### MRI brain scans

Several structural and functional brain MRI measures are available in UK Biobank as imaging derived phenotypes (IDPs)^25^. The brain imaging data, as of January 2021, were used (N=47,920). Brain imaging data used here were processed and quality-checked by UK Biobank and we made use of the IDPs^26,27^. Details of the UK Biobank imaging acquisition and processing, including structural segmentation and white matter diffusion processing, are freely available from three sources: the UK Biobank protocol: http://biobank.ctsu.ox.ac.uk/crystal/refer.cgi?id=2367 and documentation: http://biobank.ctsu.ox.ac.uk/crystal/refer.cgi?id=1977 and in protocol publications (https://biobank.ctsu.ox.ac.uk/crystal/docs/brain_mri.pdf). Total white matter hyperintensity volumes were calculated on the basis of T1 and T2 fluid-attenuated inversion recovery, derived by UK Biobank. White matter hyperintensity volumes were log-transformed due to a positively skewed distribution. We constructed general factors of white matter tract integrity using principal component analysis. The two separate unrotated factors used were fractional anisotropy (FA), gFA, and mean diffusivity (MD), gMD, previously shown to explain 54% and 58% of variance respectively^28^. We constructed a general factor of frontal lobe grey matter volume using 16 subregional volumes as per Ferguson et al^28^. Total grey matter and white matter volumes were corrected for skull size (by UK Biobank). Models were adjusted for age, sex, genetic principal components (GPCs) 1-8 and the happiness PRS. (see below)

### Adolescent Brain Cognitive Development

#### Cohort description

The Adolescent Brain Cognitive Development (ABCD) cohort is a longitudinal study of brain development and child health^10^. Investigators at 21 sites around the USA conducted repeated assessments of brain maturation in the context of social, emotional, and cognitive development, as well as a variety of health and environmental outcomes. We analysed data from release 3.0. At the time of the survey questions, the children ranged in age from 9 to 12 years. Informed written consent was provided by parents and assent was provided by children, The ABCD research protocol approved by study itself was approved by the Institutional Review Board of University of California San Diego (IRB# 160091) ^29^.

#### Genotyping

DNA was extracted from saliva samples of the ABCD participants^30^. These samples were genotyped on the Affymetrix NIDA SmokeScreen Array (Affymetrix, Santa Clara, CA, USA). The QC procedures are described in full at the following URL: https://doi.org/10.15154/1503209.

#### Phenotyping and exclusion criteria

As no direct measurement of general happiness was available, a set of questions taken from the ABCD Youth NIH Toolbox Positive Affect Items was used instead. These questions measured aspects of positive emotions and affective well-being in the past week, specifically being attentive, delighted, calm, relaxed, enthusiastic, interested, confident, energetic and able to concentrate. Responses were measured as ‘not true’, ‘somewhat true’ or ‘very true’. Each item was analysed separately.

As the initial UK Biobank GWAS was run in the white British sub-group, testing was performed firstly in the white (as defined by ABCD) participants and secondly in the whole sample, with ancestry treated as a factor variable. Participant ancestry was derived from a series of yes/no questions where the respondent’s parent selected from a list of options. Those options were: white, black, Native American, Alaskan, Hawaiian, Guamanian, Samoan, Other Pacific Islander, Indian, Chinese, Filipino, Japanese, Korean, Vietnamese, Other Asian, Other Race, Refuse To Answer, and Don’t Know. Those who responded with either of the last two options were excluded. An additional question, “Is your child Hispanic?”, was also asked and those who responded yes were defined as Hispanic irrespective of whether they also selected another option. Other categories were then defined as follows: White (n = 4,655), Black (n = 1,016), Hispanic (n = 1,201), Native American (Native American and Alaskan, n = 37), East Asian (Chinese, Filipino, Japanese, Korean, Vietnamese and Other Asian, n = 94), Indian (n = 37), Pacific Islanders (Hawaiian, Guamanian, Samoan, Other Pacific Islander, n = 8) and Other Race (n= 39).

#### MRI

Creation of the derived MRI variables from the ABCD cohort has been described in detail elsewhere^31^. For the purposes of this study, total frontal lobe volume was derived by summing the 22 frontal lobe subsection variables of the left and right hemisphere^32^. Additionally, we looked at total grey and white matter volume and left and right hippocampus volume. The hippocampal body and tail regions and white matter hyperintensity volume were not available for replication. All outcomes were transformed into z scores and all models were adjusted for age, sex, PGCs 1-8, MRI site, and the happiness PRS. For models that included participants from different ancestries, a factor variable for ancestry was included. Models were weighted to match the American community survey (ACS) data by the weighting variable “acs raked propensity score”. Relationship filtering was also performed removing one individual at random from any pair of participants who were 2^nd^ cousins or closer with valid phenotypes.

### Add Health

#### Cohort description

Add Health is a nationally representative cohort study of more than 20 000 adolescents from the USA who were aged 12–19 years at baseline assessment in 1994–95. They have been followed through adolescence and into adulthood with five in-home interviews in five waves (I-V) conducted in 1995, 1996, 2001–02, 2008–09 and 2016–18. In this analysis, participants ranged from 24.3 – 34.7 years old, 53% were female and 62% were non-Hispanic white. The study was approved by the University of California San Diego Institutional Review Board (IRB #190002XX).

#### Genotyping

Saliva samples were obtained as part of the Wave IV data collection. Two Illumina arrays were used for genotyping, with approximately 80% of the sample genotyped with the Illumina Omni1-Quad BeadChip and the remainder of the group genotyped with the Illumina Omni2.5-Quad BeadChip. After quality control, genotyped data were available for 9,974 individuals (7,917 from the Omni1 chip and 2,057 from the Omni2 chip) on 609,130 SNPs present on both genotyping arrays^33^. Imputation was performed separately for European ancestry (imputed using the HRC reference panel) and non-European ancestry samples (imputed using the 1000 Genomes Phase 3 reference panel)^34^.For more information on the genotyping and quality control procedures see the Add Health GWAS QC report online at: https://addhealth.cpc.unc.edu/wp-content/uploads/docs/user_guides/AH_GWAS_QC.pdf

#### Phenotyping and exclusion criteria

The outcome variable was collected during the at-home interview of Wave IV and was derived from the response to the question: “How often was the following true during the past seven days? You felt happy.” Responses were given as: “never or rarely”; “sometimes”; “a lot of the time”; “most of the time or all of the time”; “refused”; “don’t know”. Those who responded with the latter two options were excluded. Remaining categories were coded from “never” = 0 to “all of the time” = 3.

Ancestry in Add Health is defined in the ‘psancest’ variable as European, African, Hispanic and East Asian. Additionally, Add Health provides a weighting variable to make the results reflective of the US population. In these analyses the models were weighted by the Wave IV variable ‘gswgt4_2’. All p values for PRS analyses were False Discovery Rate (FDR)-adjusted^35^.

### Analyses

#### Genetic Association

Genetic association analysis was performed in UK Biobank using BOLT-LMM^9,36^ treating the outcome variable as a quasi-quantitative trait. BOLT-LMM uses a genetic relationship model constructed from genotyped SNPs, which allows for maximisation of sample size by adjusting for relatedness, thus also avoiding the need to adjust the model for GPCs. The model was adjusted for age, sex and genotyping array. SNPs were filtered by minor allele frequency (MAF) > 0.01, Hardy-Weinberg equilibrium p > 1 × 10^−6^, and imputation quality score > 0.8, leaving 9,286,407 SNPs for testing. BOLT-REML^37^ was also used to provide a heritability estimate and λ_GC_ estimate to test for inflation of the test statistics.

#### Linkage disequilibrium score regression

Linkage disequilibrium (LD) score regression was performed using LDSC^38^. The summary statistics from the GWAS carried out using the UK Biobank data were compared against four of the ‘big five’ personality traits: openness, conscientiousness, extraversion and agreeableness from Lo et al^39^. For neuroticism we used the summary statistics from a GWAS reported in Smith et al^40^ due to the larger sample size and a greater number of significant loci found compared to Lo et al’s^39^ neuroticism output.

### LDpred risk score generation

#### UK Biobank

LDpred^41^ established the LD structure of the genome using a reference panel of 1000 unrelated white British UK Biobank participants (the PRS discovery set). These participants had been excluded from the GWAS, due to having a missing phenotype or because they had valid MRI data, but still passed the same QC as described above. SNPs were filtered using the same parameters as for the GWAS. Scores were then created in the validation set using an infinitesimal model. Models using polygenic risk scores (PRS) derived using LDpred were adjusted for age, sex, genotyping array and the first eight GPCs.

#### ABCD and Add Health

Due to the lower cohort size of ABCD and Add Health, it would not have been possible to remove 1000 participants from the analyses to use as a discovery set without markedly reducing the power of the analyses. Therefore, we used the 1000 unrelated UK Biobank participants as the discovery set to establish LD and this was used to generate the risk scores for the participants in these datasets^42^. The only additional step was to find the SNPs that were found in both the discovery and validation datasets and passed the same SNP filtering criteria in both datasets, with an additional filter that MAF threshold was set at > 0.05 due to the lower mean imputation quality of the less common SNPs found in these smaller cohorts. Models were additionally adjusted for age at view, sex and the first 10 GPCs. For multi-ancestry models, ancestry was treated as a factor variable.

### Pruning and Threshold PRS generation in ABCD and Add Health

To ensure that using the UK Biobank participants as a reference panel was not introducing biases into the LDpred risk scores in the ABCD and Add Health cohorts, pruning and thresholding (P&T) PRSs were also created using participants from within the respective cohorts to corroborate the findings of the LDpred PRS models. Linkage disequilibrium was established using a reference of unrelated white participants from each their respective cohorts, as described above. SNPs were clumped in plink^43^ using a cut-off of r^2^ > 0.1 in a 500kb window. SNPs were filtered using p-value thresholds of p < 5 × 10^−5^, p < 0.01, p < 0.05, p < 0.1, p < 0.5 and p < 1.

